# Effects of sublethal concentrations and application concentration of SYP-9625 on *Tetranychus cinnabarinus* (Boisduval) and its natural enemy, *Neoseiulus californicus* (McGregor)

**DOI:** 10.1101/340521

**Authors:** Jingqi Ouyang, Yajing Tian, Chunxian Jiang, Qunfang Yang, Haijian Wang, Qing Li

## Abstract

**Objective:** Exploring the effects of acaricides on predatory mites is crucial for the combination of biological and chemical control of pests. In this study, sublethal effects of the new acaricide SYP-9625 on *Tetranychus cinnabarinus* (Boisduval), effects of application concentration of SYP-9625 on the predatory mite *Neoseiulus californicus* (McGregor) and functional responses of *N. californicus* were assessed. The aim of the present study was to evaluate and explore the application of new acaricide SYP-9625 with natural enemy *N. californicus*.

**Method:** All the experiments were under laboratory conditions [25 ± 1 °C, 16:8 (L:D) h and 75 ± 5% RH] and based on an age-stage, two-sex life table. The sublethal concentrations against *T. cinnabarinus*, including LC_10_ (0.375 μg/mL) and LC_30_ (0.841 μg/mL) and the application concentration (100 μg/mL) of SYP-9625, were used to evaluate effects on population parameters of *N. californicus*.

**Result:** *T. cinnabarinus* females treated with LC_30_ exhibited significantly reduced net reproductive rates (*R_0_*=11.02) of offspring compared to females treated with LC_10_ (*R_0_*=14.96) and untreated females (*>R_0_*=32.74). However, the intrinsic rate of increase (*r_m_*) and finite rate of increase (*λ*) of *N. californicus* indicated that the application concentration of SYP-9625 had no significant negative effect on treated *N. californicus* eggs (*r_m_*=0.277, *λ*=1.319) compared to the control (*r_m_*=0.292, *λ*=1.338). Additionally, the sublethal concentrations against *T. cinnabarinus* including LC_10_ and LC_30_ showed a dose-dependent mechanism on the predatory mite. SYP-9625 also stimulated the predatory capacity of *N. californicus* against immobile stages such as eggs and larvae.

**Conclusion:** It is demonstrated that sublethal concentrations of SYP-9625 can inhibit population growth of *T. cinnabarinus*. And the sublethal concentrations and application concentration had little effect on the population growth of *N. californicus*. The two advantages showed great commercial potential of this new acaricide. Therefore, *N. californicus* can manage *T. cinnabarinus* populations effectively with appropriate SYP-9625 concentrations.

## Introduction

Nowadays, agricultural spider mite pests are becoming serious threat to some important crops such as vegetables, fruits and ornamentals throughout the world. Most spider mite pests, such as *Tetranychus cinnabarinus* (Boisduval), have rapid development of resistance resulting from frequent applications of acaricides [1, 2]. Therefore, new acaricides which have excellent insecticidal activity and low toxicity to natural enemies is increasingly needed [3].

In the past few years, in order to combine natural enemies and acaricides, some studies tend to paying more attention to the toxicity of acaricides toward predatory mites. There might be inter-population differences in the sensitivities of these natural enemies [4]. Lima evaluated different acaricide toxicity against *Neoseiulus barkeri* (Hughes) and suggested that fenpyroximate and chlorfenapyr can be used with predatory mite application [5]. Recently, sublethal effects of acaricides are more considered than direct contact toxicity as an accurate approach to measure toxicity [6]. Argolo found that the sublethal concentration of imidacloprid negatively affected the population parameters of *N. californicus*, which provided a theoretical basis that the compatibility of acaricides with natural enemies can be used to manage agricultural pests [7]. Furthermore, the impact of insecticide on the functional response of the predators was considered as a way to evaluate the effect of insecticide on predation ability. The sublethal effects on powerful predation capacity of predatory mite species are worth to discuss as well. Poletti reported that acetamiprid did not affect the functional responses of *N. californicus* significantly, but it weakened the predation capacity of *Phytoseiulus macropilis* (Acari: Phytoseiidae) [4].

The predatory mite, *Neoseiulus californicus* (McGregor) (Acari: Phytoseiidae) is one of the principal natural enemies of tetranychid mites in several countries and thus promotes efficient control of those mites in several crops [8]. Moreover, *N. californicus* also exhibits broad environmental tolerance. This predatory mite species is used to manage pest mite populations in many countries, thus demonstrating the great biological control potential of *N. californicus* [9–12].

SYP-9625 is a new acaricide synthesized by Yu et al, targeting *T. cinnabarinus*, and obtained from Shenyang Sinochem Agrochemicals R&D Company Ltd. The CAS number is 1253429-01-4 [2]. SYP-9625 is one of a series of novel pyrazolyl acrylonitrile derivatives which showed excellent acaricidal activity against *T. cinnabarinus* and quite low toxicity to mammals. Its structure was identified by combination of ^1^H NMR, ^13^C NMR, and MS spectra, as well as further confirmed by X-ray diffraction.

As a consequence, this study investigated the sublethal effects of the new acaricide SYP-9625 on *T. cinnabarinus* and effects of application concentration of SYP-9625 on the predatory mite *N. californicus* after feeding on *T. cinnabarinus* based on the theory of age-stage, two-sex life tables. In this study, functional responses of *N. californicus* were also assessed to test its predation capacity. The aim of the present study was to evaluate the application of new acaricide SYP-9625 with natural enemy *N. californicus*. And try to find the reasonable concentration of SYP-9625 which has excellent insecticidal activity and low toxicity to *N. californicus*.

## Materials and methods

### Cultures

The *N. californicus* colony was originally sampled in the Sichuan Province, China in 2010 and was reared on detached kidney bean plants (*Phaseolus vulgaris* L.) infested with *T. cinnabarinus* in the laboratory. The *T. cinnabarinus* colony was collected from a farm located at Sichuan Agricultural University, China. Both laboratory colonies were maintained in the laboratory at a photoperiod of 16:8 (L: D) h, 25 ± 1°C and 75 ± 5% RH. To hold water, 9 cm diameter glass petri dishes were used to construct rearing arenas that were sealed using plastic wrap. A thin cotton layer was placed at the bottom of the petri dish, and then an upturned bean leaf was placed on the saturated cotton and surrounded with water to prevent the escape of mites. The kidney bean leaves were replaced with new ones every week. All tests were conducted at a photoperiod of 16:8 (L: D) h, 25 ± 1°C and 75 ± 5% RH in the laboratory [3].

### Chemical tested

SYP-9625 is a new acaricide synthesized (Fig 1) by Yu et al, targeting *T. cinnabarinus*, and obtained from Shenyang Sinochem Agrochemicals R&D Company Ltd. The CAS number is 1253429-01-4 [2]. SYP-9625 is one of a series of novel pyrazolyl acrylonitrile derivatives in an international patent named a pyrazolyl acrylonitrile compounds and uses thereof [13]. Application number is WO2010CN72224 20100427 and Priority number is CN2009183205 20090429. According to the patent, a pyrazolyl acrylonitrile compounds represented by the structures of Formula I: or stereoisomers thereof, wherein: R_1_ is selected from the group of substituents consisting of H, C_1_-C_4_ alkoxy C_1_-C_2_ alkyl, C_3_-C_5_ alkenyloxy C_1_-C_2_ alkyl, C_3_-C_5_ alkynyloxy C_1_-C_2_ alkyl, C_1_-C_4_ alkylthio C_1_-C_2_ alkyl, C_1_-C_5_ alkyl carbonyl, C_3_-C_8_ cycloalkyl carbonyl, C_1_-C_5_ alkoxy carbonyl or C_1_-C_5_ alkylthio carbonyl; R_2_ is C_l_ or methyl; R_3_ is H, methyl, CN, NO_2_ or halogen. Yu investigated the syntheses and bioactivities of SYP-9625, and showed its excellent acaricidal activity against *T. cinnabarinus* and low acute toxicity to mammals.

**Fig 1.**
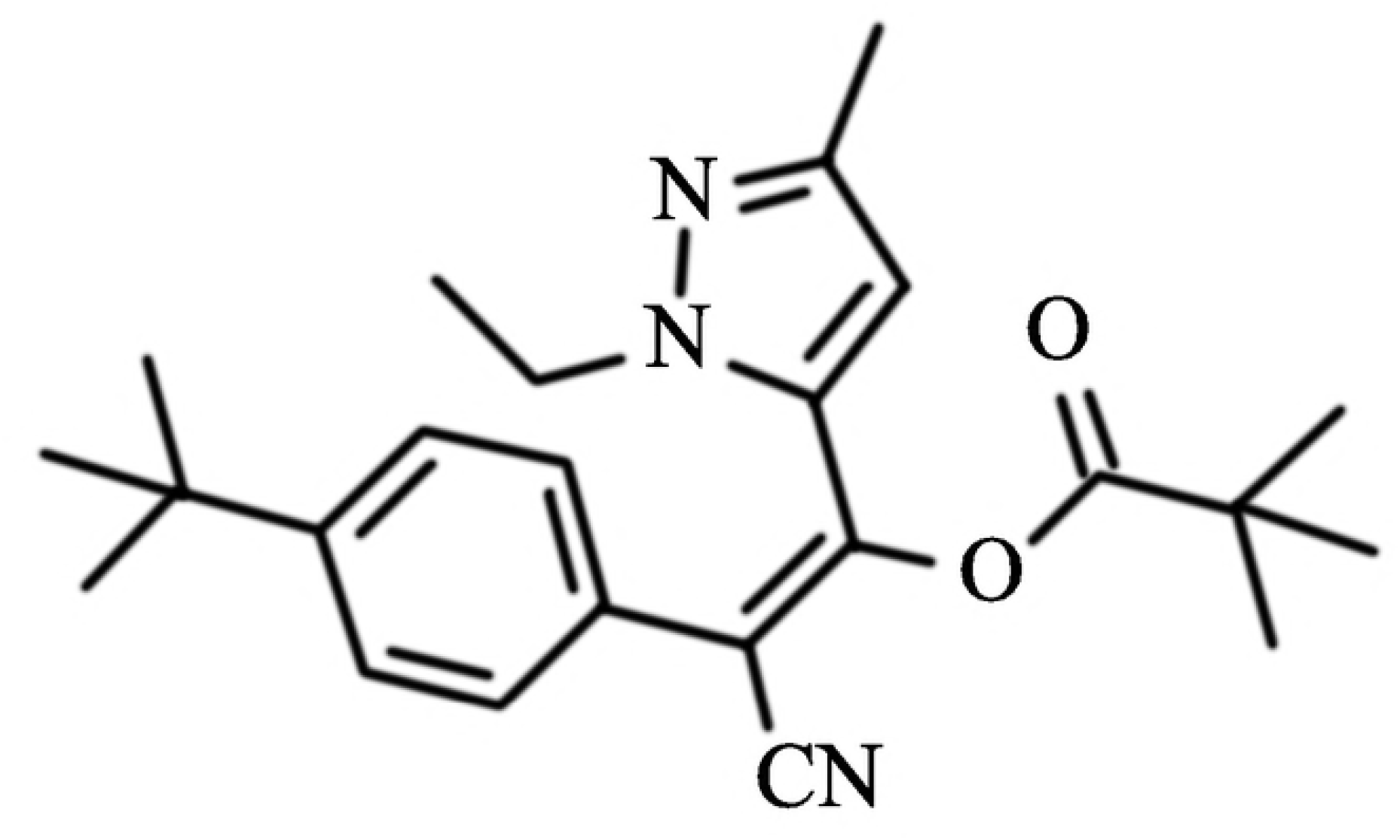
Structure of SYP-9625

### Selection of sublethal concentrations of SYP-9625

A modified Leaf-residue method was used to the response of *T. cinnabarinus* to various concentrations of SYP-9625. Bean leaf disks (2 cm diameter) were immersed for 5 s in the solutions of SYP-9625 which were chosen based on initial rang-finding tests and 0.05% Tween 80 aqueous solution (control) and allowed to dry. Then, we transferred healthy *T. cinnabarinus* females onto the bean leaf disks after their eclosions. After 24 h, mites were separated onto untreated leaf disks to couple with males from the stock colony. Every 12 h, the fecundity of females was recorded until the females naturally died [3]. There were 30 individuals per replicate and four replicates per concentration.

A modified leaf dip method [14] was used to the response of *T. cinnabarinus* eggs to various concentrations of SYP-9625. A 2 cm diameter leaf disk with 30 pregnant *N. californicus* adults was placed in a petri dish that was supported by a cotton pad filled with distilled water. After 12 h, mite females were placed on leaves to oviposit and then removed. The bean leaves with 50 eggs were dipped for 5 s in the the solutions of SYP-9625 which were chosen based on initial rang-finding tests and 0.05% Tween 80 aqueous solution (control) and then placed upside down on a wet cotton pad filled with distilled water. Eggs were checked every day and hatched in the laboratory. There were four replicates per concentration.

The two methods were also used for assessing the response of *N. californicus* females and eggs to the application concentration of SYP-9625 (100 μg/mL) and even ten times of it (100 μg/mL).

### Experimental set up

**To assess the effects of the sublethal concentration of SYP-9625 on *T. cinnabarinus* and its offspring**, bean leaf disks (2 cm diameter) were immersed for 10 s in sublethal concentrations (LC_10_ and LC_30_) and 0.05% Tween 80 aqueous solution (control) and allowed to dry. Then, we transferred healthy females of *T. cinnabarinus* onto the treated and untreated bean leaf disks after their eclosions. After 24 h, mites were separated onto untreated leaf disks to couple with males from the stock colony. Every 12 h, the fecundity of females was recorded until the females naturally died. Approximately 100 to 120 eggs were saved and transferred onto untreated bean leaf disks. The population parameters were recorded every 12 h after eclosions of both sexes. The offspring were coupled with males from the stock colony and all indices were recorded until females naturally died [3, 15].

**To assess the effects of the application concentration on *N. californicus* eggs**, a 3.5 cm diameter leaf disk with enough *T. cinnabarinus* at every life stage, as well as with thirty pregnant *N. californicus* adults, was placed in a petri dish that was supported by a cotton pad filled with distilled water. After 12 h, mite females were placed on leaves to oviposit and then removed. The bean leaves with 35 eggs were dipped in the solution of application concentration (100 μg/mL) for 5 s per disk and then placed upside down on a wet cotton pad filled with distilled water. Eggs were checked every day and hatched in the laboratory. After hatching, they were separated onto the untreated 2 cm diameter leaf disks using 0.05% Tween −80 aqueous solution as a control. Population parameters were recorded every 12 h after their eclosions, considering both sexes, and all indices were recorded until all females died [16].

**We also tested the effects of the application concentration on *N. californicus* and its offspring from treated females**. The treatment method and device were similar to the experiment about sublethal effect of concentrations on *T. cinnabarinus* and its offspring from treated females.

**A modified method was considered to assess the indirect effect on *N. californicus* and its offspring fed on sublethal treated *T. cinnabarinus***. We fed *N. californicus* on treated females of *T. cinnabarinus* and evaluated population parameters of predator mites. Enough eggs of *N. californicus* fed on untreated *T. cinnabarinus* were collected in 24 h. When *N. californicus* grew to the deutonymph life stage, enough *T. cinnabarinus* females were treated at sublethal concentrations (LC_10_ and LC_30_) and 0.05% Tween-80 aqueous solution (control) using the same method as described above. After 24 h, the *N. californicus* were fed on the treated *T. cinnabarinus* females and coupled with males from the stock colony. Then, population parameters were recorded every 12 h considering both sexes and all indices were recorded until females died.

There were 60 individuals of *N. californicus* per replicate and four replicates per concentration.

**To assess the effects of application concentration on the predation functional response of *N. californicus***, bean leaf disks (4 cm diameter) were immersed in the application concentration (100 μg/mL) and 0.05% Tween-80 aqueous solution (control) and allowed to dry. Healthy *N. californicus* females were transferred onto the treated and untreated bean leaf disks within 12 h of copulation. After 24 h, they were separately transferred onto untreated bean leaf disks (1 cm×0.5 cm) and fed with *T. cinnabarinus* at every stage. Eggs and nymphs were transferred for 10, 15, 20, 25, and 30 per leaf. Larvae were transferred for 10, 20, 30, 40, and 50 per leaf. Adults were transferred for 10, 15, 20, 25, and 30 per leaf. All leaves were placed in centrifuge tubes (2 ml), which were specially made to prevent mites from escaping [17].

**To assess the effects on functional response of *N. californicus* fed on sublethal treated *T. cinnabarinus***, healthy *N. californicus* females were separately introduced onto freshly cut leaf disks (2 mm×5 mm) that were placed in centrifuge tubes (0.5 ml) 12 h after copulation for host starvation for 24 h. *T. cinnabarinus* on every stage were treated for 24 h by sublethal concentrations (LC_10_ and LC_30_) and 0.05% Tween −80 aqueous solution (control) using the same method as the effect of sublethal concentrations on *T. cinnabarinus* and its offspring from treated females. *T cinnabarinus* were transferred onto leaf disks (1 cm×0.5 cm) with separately treated *N. californicus*, and the density setting was same as described above (see Effect of application concentration on the functional response of *N. californicus* from treated females).

There were five replicates per concentration. The functional response of *N. californicus* was observed and recorded after 24 h.

### Statistical analysis

The means and standard errors of the population parameters were estimated using a paired bootstrap test (TWOSEX-MS Chart) procedure [18] because it uses random resampling. Few replications can generate variable means and large standard errors (P<0.05); thus, we used 10,000 replications.

The functional responses of *N. californicus* to the various prey stages and densities were expressed by fitting Holling’s equation to the data [19, 20]:

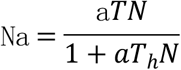

Where Na is the number of prey eaten, T is the experimental time (1h), *N* is the initial number of prey offered, a is the searching (attack) rate, and *T_h_*is the handling time. Searching rate, handling time and their asymptotic standard errors were estimated from nonlinear regressions of the disk equation. SAS statistical software was used to analyze functional responses of *N. californicus*.

### Age-stage, two-sex life table

It is widely accepted that age-stage, two-sex life table was developed where both males and females are included to express the age-stage-structure of a population more accurately. The raw data of the life table parameters were assessed on the basis of the theory of an age-stage, two-sex life table [21–26] by the computer program TWOSEX-MS Chart [27]. The age-stage specific survival rate (*s_xj_*) (where *x*=age in days and *j*=stage), female age-specific fecundity (*f*_*x*5_), the age-specific survival rate (*l_x_*), the age-specific fecundity (*m_x_*), *m_x_* in total population and age-specific maternity (*l_x_m_x_*) and the population growth parameters [the intrinsic rate of increase (*r_m_*), the finite rate of increase (*λ*), the net reproductive rate (*R_0_*), the gross reproductive rate (GRR), and the mean generation time (T) and the doubling time (DT)] were calculated accordingly [3, 28].

## Results

### Determination of sublethal concentrations

It is revealed that SYP-9625 was selected as a candidate for extensive laboratory bioassays and field trials [2]. Moreover, according to our result of acaricidal activities of sublethal concentrations of SYP-9625, it showed excellent activity against all developmental stages of *T. cinnabarinus*. The LC_50_ of SYP-9625 on female adults and eggs are 0.466 *μg/mL* and 1.472μg/mL, respectively. The regression equation of concentration-mortality for females was Y=1.447+4.365X, [Y=mortality (probit), X= the log10 of concentration] (Table 1). No mortalities were recorded in the controls.

**Table 1.** The toxicity of SYP-9625 on different stages of *T. cinnabarinus*

| Stage | Method | LC-P line<br>(y=) | Correlation<br>coefficient<br>(r) | $x^2$ | df | LC<sub>50</sub>(µg/mL)95%CI |
| --- | --- | --- | --- | --- | --- | --- |
| Female | Leaf-residue<br>method | 1.447+4.365x | 0.9945 | 3.219 | 7 | 0.466<br>(0.442-0.492) |
| Nymph | Leaf-residue<br>method | 5.311+10.994x | 0.9975 | 0.537 | 5 | 0.329<br>(0.316-0.341) |
| **Larva** | Leaf-residue method | 2.003+4.746x | 0.9930 | 2.372 | 5 | 0.378<br>(0.357-0.404) |
| **Egg** | Leaf-dip method | -0.362+2.157x | 0.9950 | 1.244 | 3 | 1.472<br>(1.272-1.694) |

After treated in 24h, 48h and 72h, *N. californicus* females and eggs were both in sensitive with the application concentration of SYP-9625. Even at ten times of the application concentration, the hatching rate was 99.33±0.67. As a consequence, 100μg/mL of SYP-9625 is considered as the application concentration on *N. californicus* (Table 2).

**Table 2.** Effect of SYP-9625 on survival rate of eggs and female adults of *N. californicus*

| Acaricide | Dose μg/mL | Hatching rate (%) | Survival rate (%) | | |
| --- | --- | --- | --- | --- | --- |
|  |  |  | 24h | 48h | 72h |
| (SYP-9625) | 100 | 100.00±0.00<sup>a</sup> | 100.00±0.00<sup>a</sup> | 100.00±0.00<sup>a</sup> | 99.67±0.33<sup>a</sup> |
| | 1000 | 99.33±0.67<sup>a</sup> | 100.00±0.00<sup>a</sup> | 100.00±0.00<sup>a</sup> | 99.00±0.58<sup>a</sup> |

Note: Data in the table are mean ± SE. Data in the same group followed by different letters indicate significant difference at $P < 0.05$ Level by Duncan's new multiple ranges test.

The sublethal concentrations including LC_10_ (0.375μg/mL) and LC_30_ (0.841μg/mL) were determined using a probit procedure (SAS Institute 2002) for the subsequent experiments and summarized in Table 3.

**Table 3.** Sublethal concentrations of SYP-9625 on different stages of *T. cinnabarinus*

| Stage | Method | LC<sub>10</sub>(μg/mL) 95%CI | LC<sub>30</sub> (μg/mL) 95%CI |
| --- | --- | --- | --- |
| **Female** | Leaf-residue method | 0.237<br>(0.216~0.256) | 0.353<br>(0.332~0.374) |
| **Nymph** | Leaf-residue method | 0.251<br>(0.232~0.267) | 0.295<br>(0.279~0.307) |
| **Larva** | Leaf-residue method | 0.203<br>(0.182~0.221) | 0.293<br>(0.275~0.331) |
| **Egg** | Leaf-dip method | 0.375<br>(0.269~0.480) | 0.841<br>(0.688~0.990) |

### Effects of sublethal concentrations of SYP-9625 on *T. cinnabarinus* females and their offspring

Total spawning rate, female longevity and fecundity per female treated with the sublethal concentrations (LC_10_, LC_30_) on *T. cinnabarinus* were significantly reduced and the pre-oviposition periods were significantly extended compared to the controls (Table 4). Oviposition period of females treated with LC_30_ was significantly shorter than oviposition period of females in the control treatment. Total spawning rate, female longevity and fecundity per female treated with LC_30_ were lower than the LC_10_ treatment. Moreover, the pre-oviposition periods treated with LC_30_ were longer than the LC_10_ treatment.

**Table 4.**
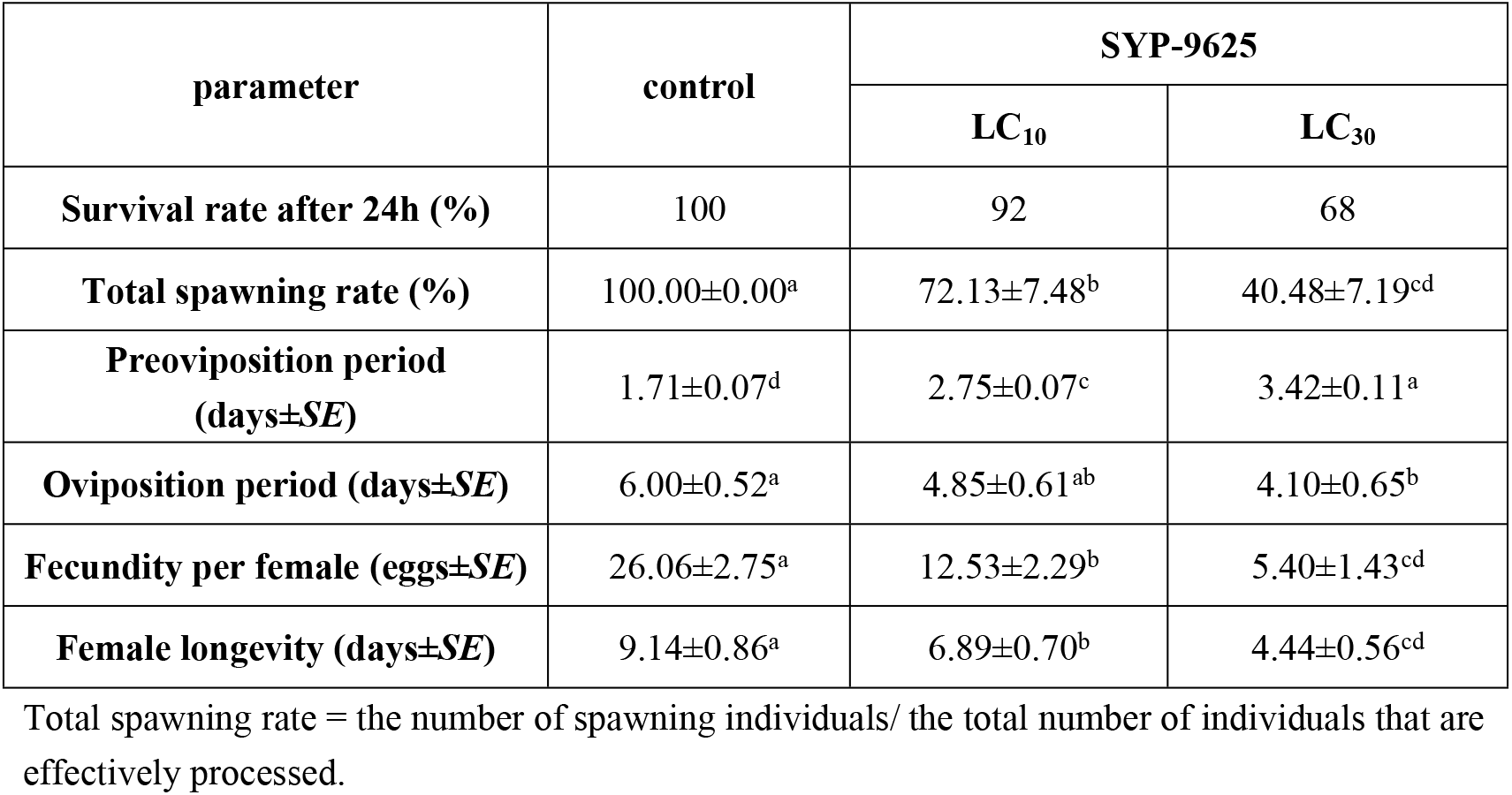
Effects of sublethal exposure of SYP-9625 on fecundity and longevity of the treated females of *Tetranychus cinnabarinus*

| parameter | control | SYP-9625 |  |
| --- | --- | --- | --- |
|  |  | LC<sub>10</sub> | LC<sub>30</sub> |
| Survival rate after 24h (%) | 100 | 92 | 68 |
| Total spawning rate (%) | 100.00±0.00<sup>a</sup> | 72.13±7.48<sup>b</sup> | 40.48±7.19<sup>cd</sup> |
| Preoviposition period (days±SE) | 1.71±0.07<sup>d</sup> | 2.75±0.07<sup>c</sup> | 3.42±0.11<sup>a</sup> |
| Oviposition period (days±SE) | 6.00±0.52<sup>a</sup> | 4.85±0.61<sup>ab</sup> | 4.10±0.65<sup>b</sup> |
| Fecundity per female (eggs±SE) | 26.06±2.75<sup>a</sup> | 12.53±2.29<sup>b</sup> | 5.40±1.43<sup>cd</sup> |
| Female longevity (days±SE) | 9.14±0.86<sup>a</sup> | 6.89±0.70<sup>b</sup> | 4.44±0.56<sup>cd</sup> |

Total spawning rate = the number of spawning individuals/ the total number of individuals that are effectively processed.

Fig 2 illustrates that age-specific fecundity curves and peak values of *T. cinnabarinus* adult females treated with sublethal concentrations (LC_10_, LC_30_) moved backward Moreover, a significant reduction of age-specific survival rate was observed with different concentrations.

**Fig 2.**
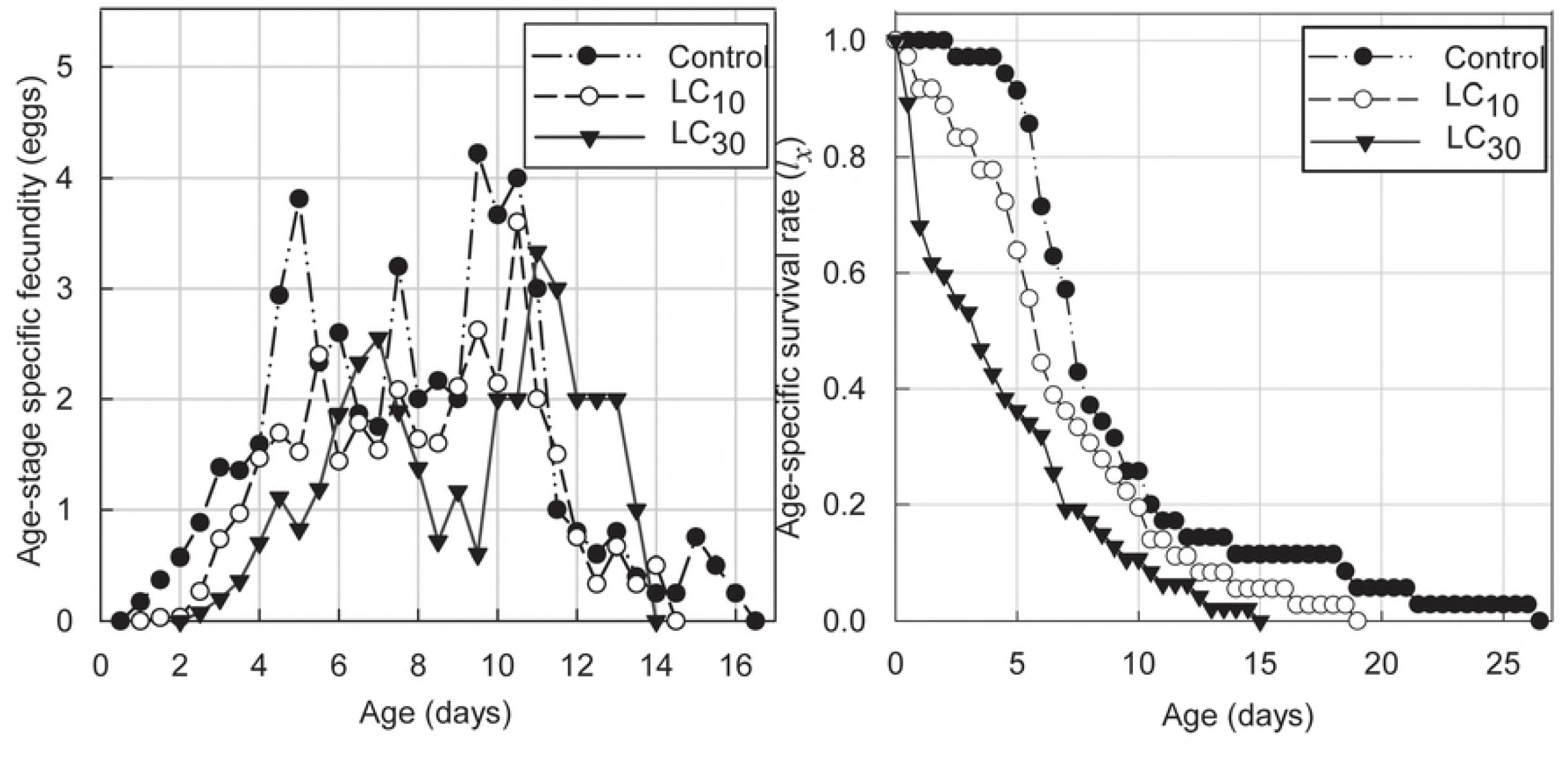
Female age-specific fecundity (*fx*_5_), Age-specific survival rate (*lx*) of *T. cinnabarinus* female adults treated with sublethal concentrations of SYP-9625. After assessed effects of sublethal concentrations on offspring from treated *T. cinnabarinus* females, the growth rates in pre-adult durations at LC_10_ and LC_30_ were lower than in the control. The curves moved backward and there was less recombination (Fig 3). Moreover, the total survival rate at LC_30_ was lower than the LC_10_ treatment and the control. In Fig 4, the slope of *l_x_* became steeper after 5 to 16 days as sublethal concentration ranged from LC_10_ to LC_30_, but they converged on the same value. The peak values of *f*_*x*5_, *l_x_m_x_* in individuals who survived treatments LC_10_ and LC_30_ were distinctly lower than the control, but less difference was observed between LC_10_ and LC_30_. Consequently, sublethal concentrations of SYP-9625 weakened the population reproductivity, especially the fecundity of female mites.

**Fig 3.**
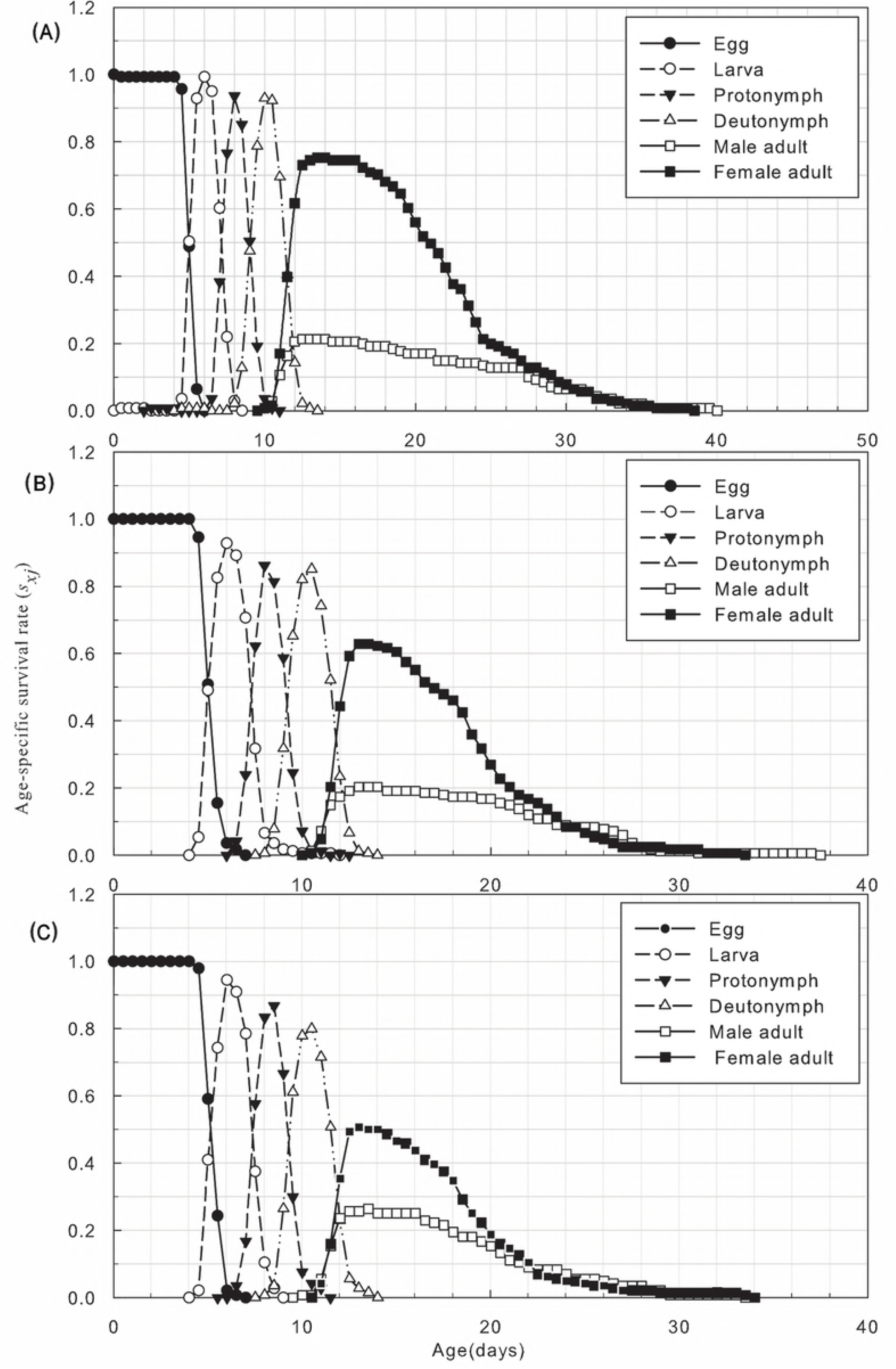
Age-stage specific survival rate (*s_xj_*) of offspring from females of *T. cinnabarinus* treated with sublethal concentrations of SYP-9625. (A) Control, (B) LC10, (C) LC30.

**Fig 4.**
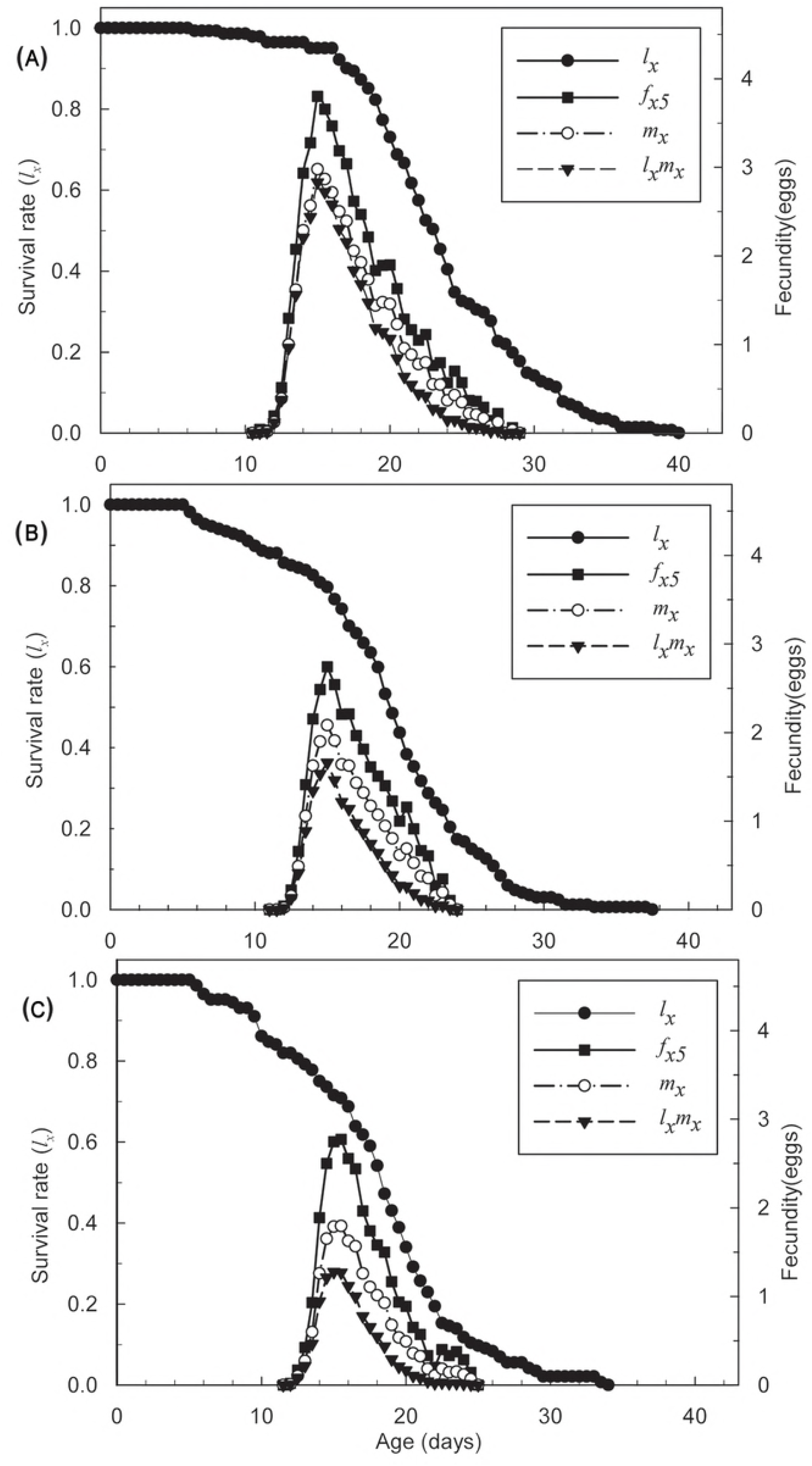
Age-specific survival rate (*l_x_*), female age-specific fecundity (*f*_*x*5_), age-specific fecundity of total population (*m_x_*), and age-specific maternity (*l_x_m_x_*) of *T. cinnabarinus* eggs treated with sublethal concentrations of SYP-9625. (A) Control, (B) LC_10_, (C) LC_30_.

In Table 5, *r_m_, λ* and *R_0_* of offspring from the treated *T. cinnabarinus* females were significantly lower than the control. The increasing concentration caused the dramatic change. Additionally, *T* at LC_30_ was significantly shorter than the control.

**Table 5.**
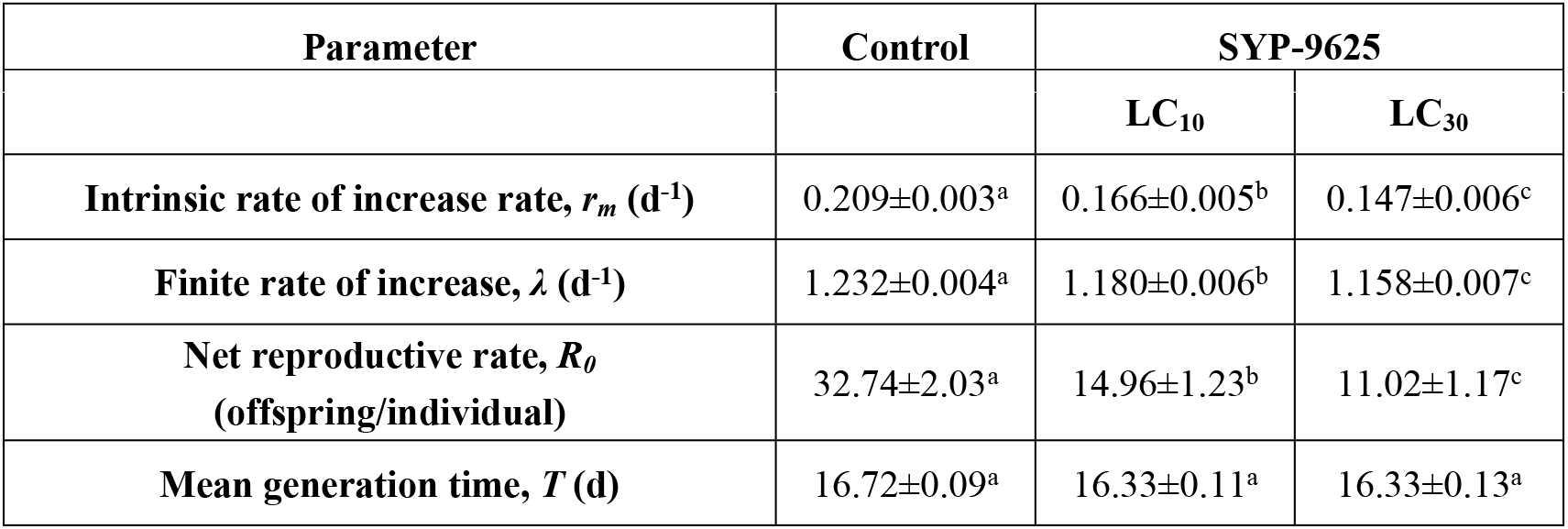
Population life table parameters of offspring from females of *Tetranychus cinnabarinus* treated with sublethal concentrations of SYP-9625

### Effects of the application concentration of SYP-9625 on *N. californicus* eggs

After a 5 seconds exposure to the application concentration (100μg/mL), preadult duration, longevity and total life span of the treated eggs of *N. californicus* were not varied in table 6. Larva and protonymph duration of treatment were longer than the control, beyond that, other indexes including female proportion and adult emergence rate had less difference with the control. Table 7 presented the similarity of spawning rate, pre-oviposition and fecundity per female among the females that grown from treated eggs. The total pre-oviposition treated with SYP-962 were significantly longer than the control, in contrast, the oviposition were shorter.

**Table 6.** Development time, longevity, and total life span of *Neoseiulus californicus* eggs treated with the application concentration of SYP-9625

| Parameter | Control | SYP-9625(100µg/mL) |
| --- | --- | --- |
| Female proportion (%) | 62.88±4.91<sup>a</sup> | 60.99±4.87<sup>a</sup> |
| Adult emergence rate (%) | 97.00±1.70<sup>ab</sup> | 100.00±0.00<sup>a</sup> |
| **Female** |  |  |
| Egg duration(d) | 1.83±0.07<sup>a</sup> | 1.84±0.03<sup>a</sup> |
| Larva duration(d) | 0.58±0.03<sup>b</sup> | 0.73 ± 0.03<sup>a</sup> |
| Protonymph duration(d) | 0.98± 0.01<sup>b</sup> | 1.05 ±0.02<sup>a</sup> |
| Deutonymph duration(d) | 1.22±0.06<sup>a</sup> | 1.27±0.25<sup>a</sup> |
| Preadult duration(d) | 4.61±0.04<sup>b</sup> | 4.89± 0.04<sup>a</sup> |
| Longevity(d) | 30.95±1.19<sup>a</sup> | 26.30±1.37<sup>b</sup> |
| Total life span (d) | 35.56±1.21<sup>a</sup> | 31.18±1.38<sup>b</sup> |
| **Male** |  |  |
| Egg duration(d) | 1.86± 0.04<sup>a</sup> | 1.91± 0.03<sup>a</sup> |
| **Larva duration(d)** | 0.58± 0.03<sup>a</sup> | 0.59± 0.03<sup>a</sup> |
| **Protonymph duration(d)** | 0.88±0.04<sup>b</sup> | 0.91±0.03<sup>b</sup> |
| **Deutonymph duration(d)** | 1.01±0.03<sup>a</sup> | 1.08±0.03<sup>a</sup> |
| **Preadult duration (d)** | 4.33±0.07<sup>b</sup> | 4.49±0.07<sup>ab</sup> |
| **Longevity(d)** | 33.04±2.22<sup>a</sup> | 29.58±2.06<sup>a</sup> |
| **Total life span (d)** | 37.38±2.25<sup>a</sup> | 34.06±2.05<sup>a</sup> |

**Table 7.** The reproductive period and fecundity of *Neoseiulus californicus* eggs treated with the application concentration of SYP-9625

| **Parameter** | **Control** | **SYP-9625(100µg/mL)** |
| --- | --- | --- |
| **Spawning rate (%)** | 100.00±0.00<sup>a</sup> | 100.00±0.00<sup>a</sup> |
| **Pre-oviposition (d)** | 1.68±0.04<sup>a</sup> | 1.72±0.05<sup>a</sup> |
| **Total pre-oviposition (d)** | 6.29±0.07<sup>b</sup> | 6.61±0.08<sup>a</sup> |
| **Oviposition (d)** | 15.77±0.44<sup>a</sup> | 13.30±0.45<sup>b</sup> |
| **Fecundity per female (eggs)** | 46.72±1.35<sup>a</sup> | 40.26±1.42<sup>ab</sup> |

The age-specific survival rate of females and age-stage specific survival rate of treatment and control were observed (Fig 5). We can barely distinguish the difference in *l_x_, f*_*x*5_ and *m_x_* in the total population between treatments and control. The peak value of *f*_*x*5_ at control was 2 showed in 11 days and the peak value of *f*_*x*5_ at application concentration was 1.8 showed in 10 days. *r_m_, λ*, GRR and *T* in treated *N. californicus* eggs were similar with the control (Table 8). Hence, there was little effect on the population growth of *N. californicus* eggs exposed to the application concentration of SYP-9625.

**Fig 5.**
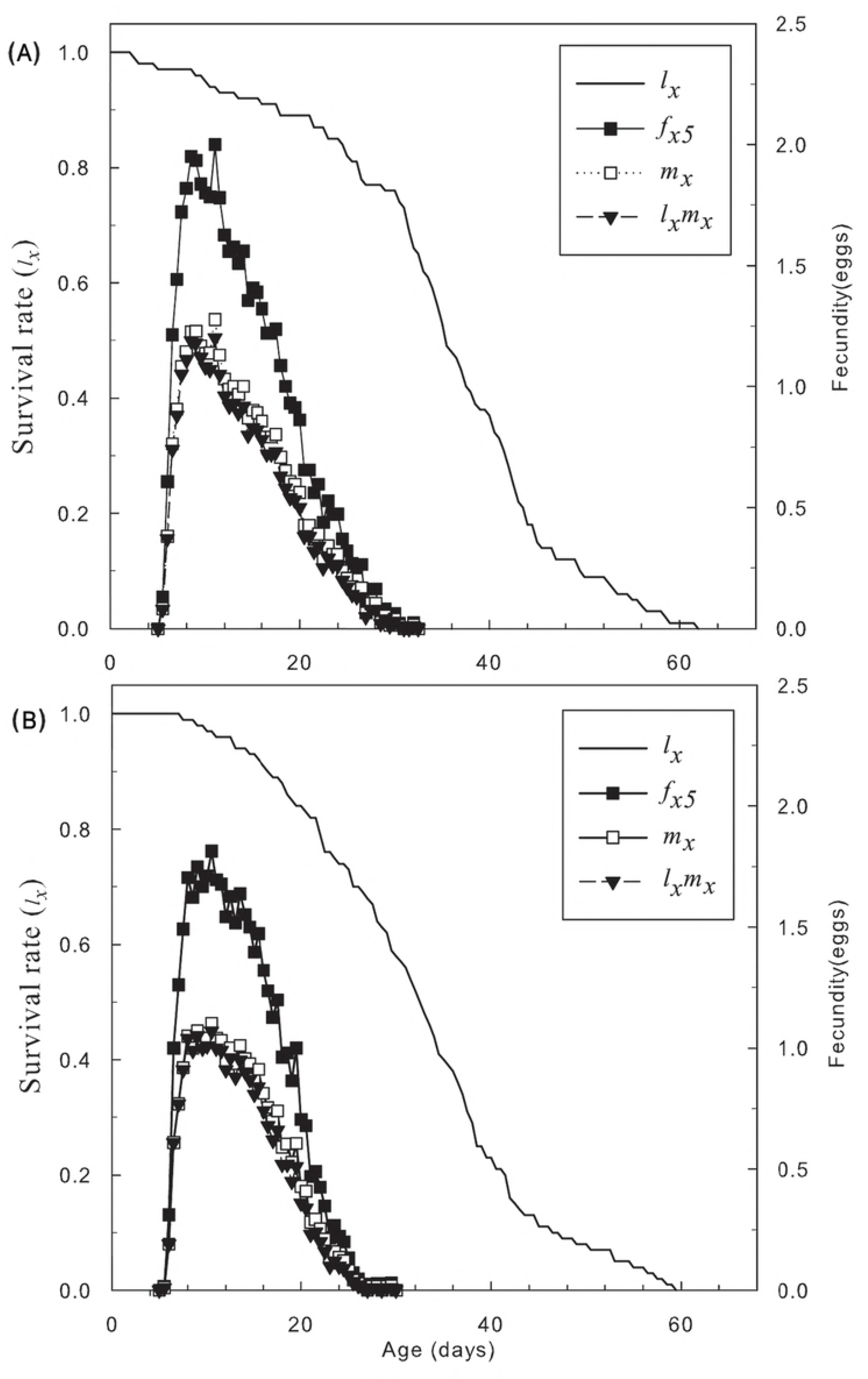
Age-specific survival rate (*l_x_*), female age-specific fecundity (*f*_*x*5_), age-specific fecundity of total population (*m_x_*), and age-specific maternity (*l_x_m_x_*) of *N. Californicus* McGregor) eggs treated with sublethal concentrations of SYP-9625. (A) Control, (B) SYP-9625.

**Table 8.** Population life table parameters of *Neoseiulus californicus* eggs treated with the application concentration of SYP-9625

| Parameter | Control | SYP-9625(100µg/mL) |
| --- | --- | --- |
| Intrinsic rate of increase rate, $r_m$ (d<sup>-1</sup>) | 0.292±0.009<sup>a</sup> | 0.277±0.009<sup>a</sup> |
| Finite rate of increase, $\lambda$ (d<sup>-1</sup>) | 1.338±0.012<sup>a</sup> | 1.319±0.012<sup>a</sup> |
| Net reproductive rate, $R_0$<br>(offspring/individual) | 28.50±2.41<sup>a</sup> | 24.56±2.15<sup>ab</sup> |
| Mean generation time, $T$ (d) | 11.49±0.14<sup>a</sup> | 11.55±0.16<sup>a</sup> |

### Effects of the application concentration of SYP-9625 on *N. californicus* females and their offspring

Fig 6 illustrates that the application concentration reduced the survival rate of treated females. The female age-specific fecundity peak value of the control was presented earlier than the treatment. Additionally, the fluctuation of female age-specific fecundity was larger than the control.

**Fig 6.**
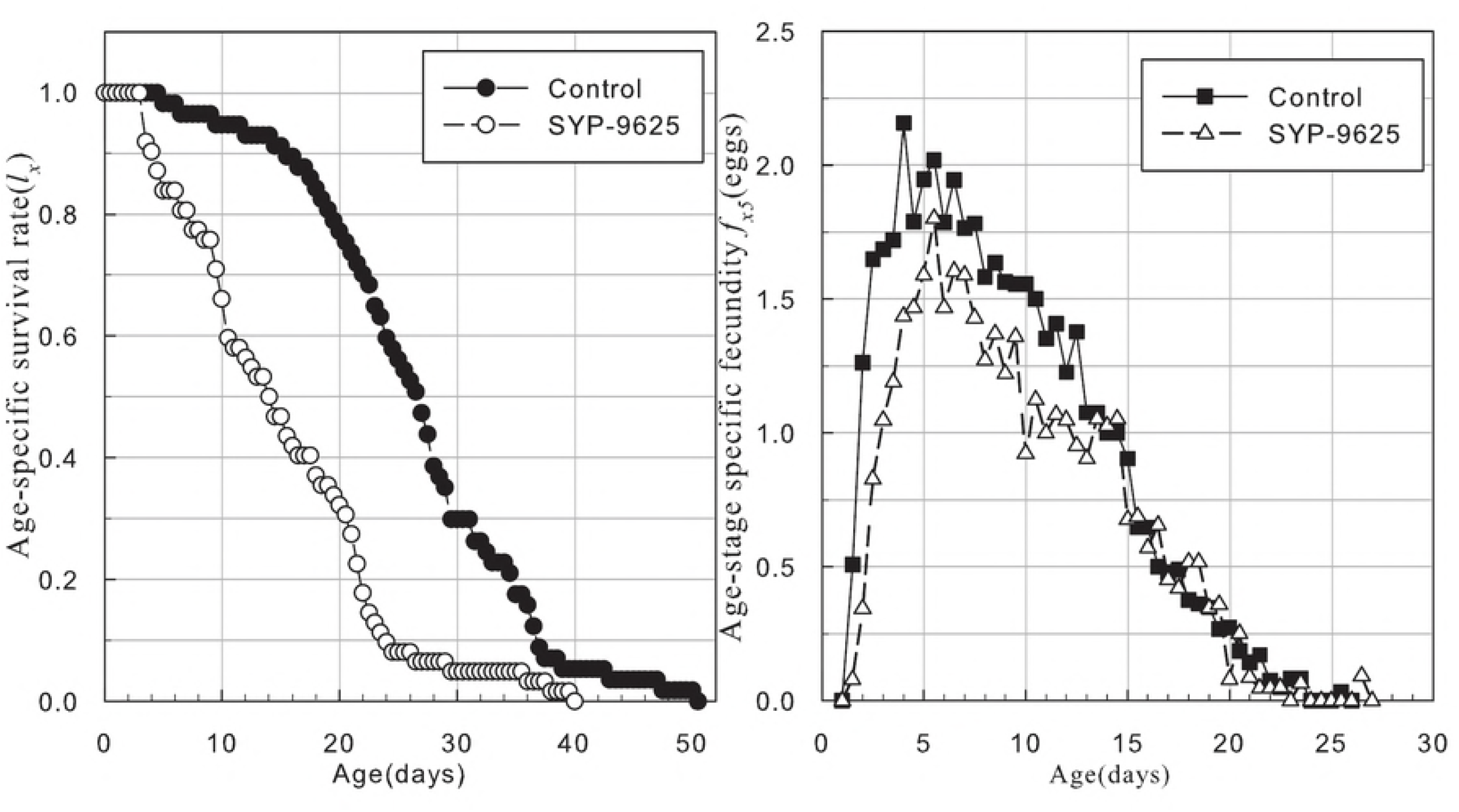
Age-specific survival rate (*l_x_*), female age-specific fecundity (*f*_*x*5_), of *N. californicus* (McGregor) female adults treated with sublethal concentrations of SYP-9625.

After assessed the effects of the application concentration on offspring from the treated *N. californicus* females, the age-stage specific survival rate of egg and protonymph ranged from 2 d to 3 d, and 3.5 d to 5 d after being treated (Fig 7). The peak value of larva survival rate was above 0.8%, which was 0.3% higher than the control. The survival rate of female adults was higher than males prior to the application concentration application, but then the survival rate of female adults decreased. The peak value was 0.4%, which was 0.2% lower than the control.

**Fig 7.**
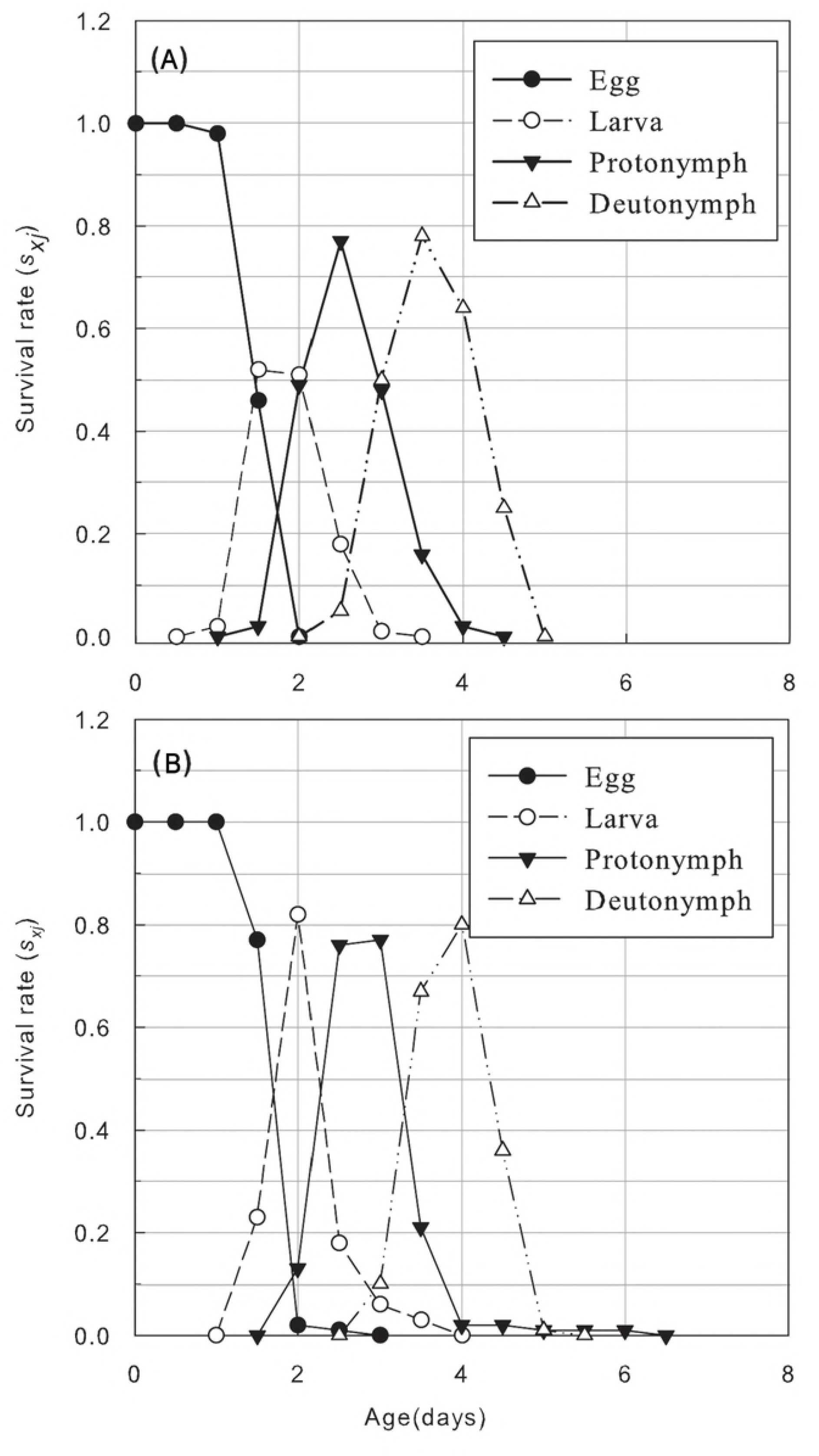
Age-stage specific survival rate (*s_xj_*) of offspring from *N.Californicus* (McGregor) female adult treated with sublethal concentrations of SYP-9625. (A)Control, (B) SYP-9625.

Initially, the age-specific survival rate at the application concentration declined slowly from 0 d to 30 d. Then, it decreased more rapidly from 30 d to 60 d, but the overall change was very minor. The acaricide treatment barely changed the age-specific survival rate of offspring from the treated females of *N. californicus*, but the peak value appeared to be delayed, and the declining gradient of the earlier stage was higher than the control (Fig 8).

**Fig 8.**
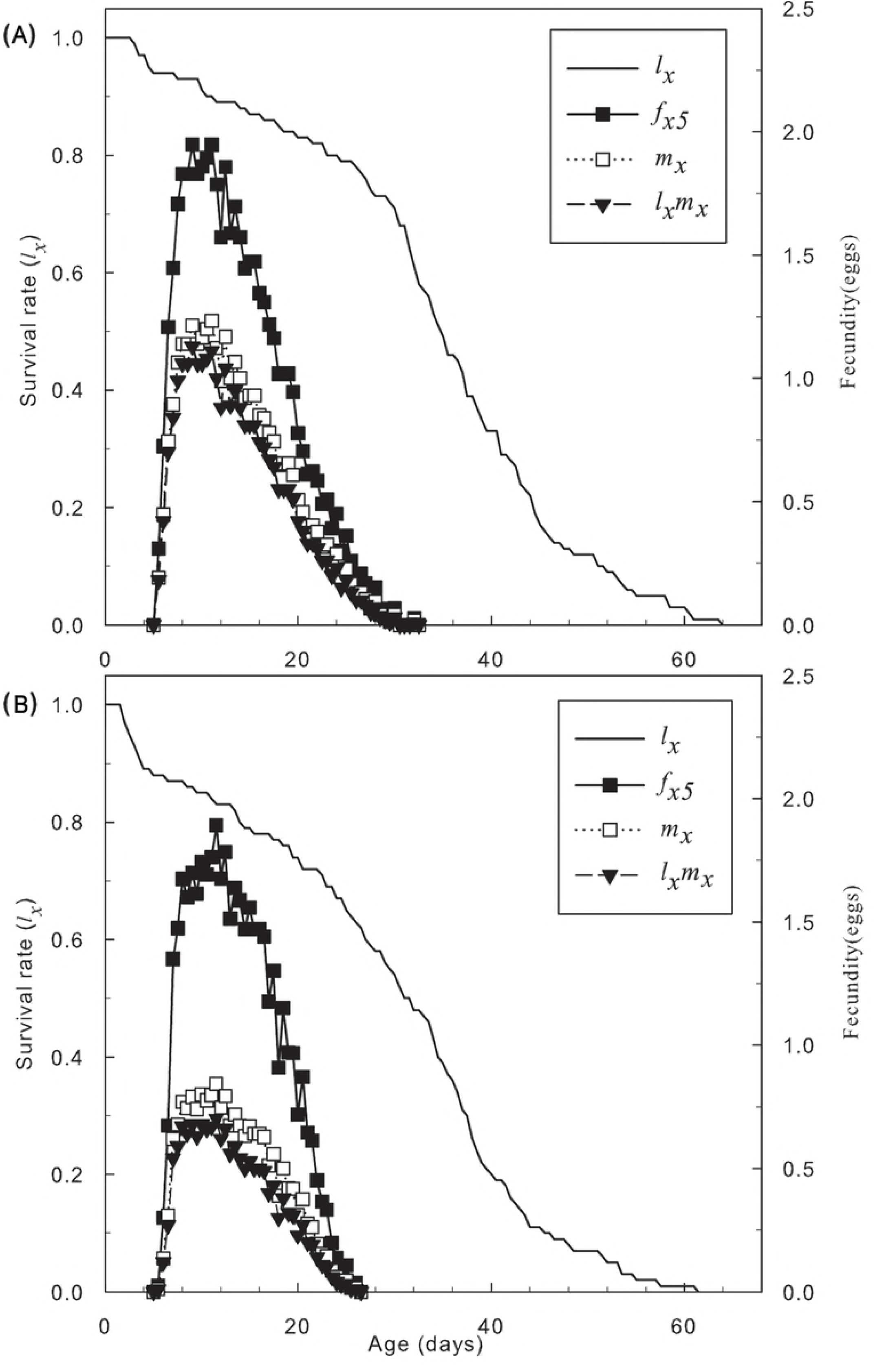
Age-specific survival rate (*l_x_*), female age-specific fecundity (*f*_*x*5_), age-specific fecundity of total population (*m_x_*), and age-specific maternity (*l_x_m_x_*) of offspring from *N.Californicus* (McGregor) female adult treated with sublethal concentrations of SYP-9625. (A) Control, (B) SYP-9625.

Lower *R_0_, r_m_* and *λ* in offspring from the treated *N. californicus* individual females treated by SYP-9625 are compared with the control in Table 9. Moreover, there was no difference of *T* observed between treatments.

**Table 9.** Population life table parameters of offspring from *Neoseiulus californicus* female treated with the the application concentration of SYP-9625

| Parameter | Control | SYP-9625(100µg/mL) |
| --- | --- | --- |
| Intrinsic rate of increase rate, $r_m$ (d<sup>-1</sup>) | 0.290±0.009<sup>a</sup> | 0.233±0.012<sup>b</sup> |
| Finite rate of increase, $\lambda$ (d<sup>-1</sup>) | 1.336±0.012<sup>a</sup> | 1.263±0.155<sup>b</sup> |
| Net reproductive rate, $R_0$<br>(offspring/individual) | 27.37±2.43<sup>a</sup> | 15.91±2.08<sup>b</sup> |
| Mean generation time, $T$ (d) | 11.42±0.14<sup>a</sup> | 11.87±0.19<sup>a</sup> |

### Effects of SYP-9625 on *N. californicus* fed on sublethal treated *T. cinnabarinus*

Fig 9 shows minor changes in the *s_xj_* of egg, larva and nymph in a dose-dependent manner. The curves demonstrate that sublethal concentrations of SYP-9625 reduced the survival of female adults, and the dose-dependent manner also emerged in the survival of male adults at the egg stage. As shown in Fig 10, *l_x_* rapidly reduced from 0 to 30 d with increased concentrations of SYP-9625, *l_x_* reduced more slowly from 30 to 48 d. After 48 d, all *l_x_* gradually decreased to 0% between 64.5 to 72.5 d. *fx*_5_, *m_x_* and *l_x_m_x_* at LC_10_ had slight differences with the control, but *fx*_5_, *m_x_* and *l_x_m_x_* at LC_30_ were all lower than the control. All the population life table parameters at LC_30_ were lower than the control with the exception of *T* (Table 10). After *N. californicus* fed on treated *T. cinnabarinus, r_m_* of its subsequent generation significantly reduced from 0.289 to 0.243. Intrinsic rate of increase rate (*r_m_*) was an important measure of variation of the population trend in the specific environment conditions to reflect the reproduction of *N. californicus*. And *λ* significantly reduced from 1.335 to 1.275, *R_0_* significantly reduced from 28.71 to 18.13. Moreover, population life table parameters at LC_10_ were appropriately similar to the control.

**Fig 9.**
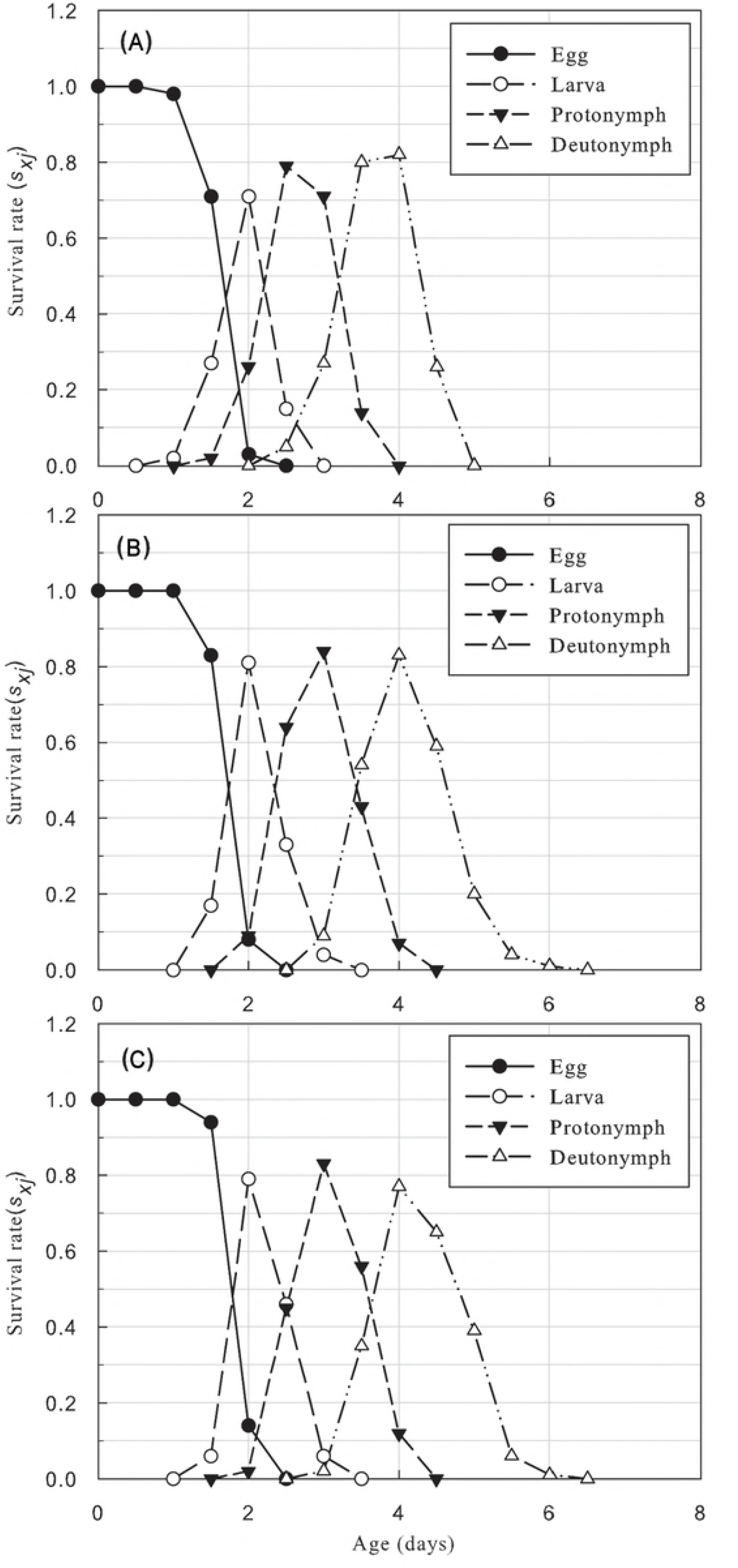
Age-stage specific survival rate (*s_xj_*) of offspring from *N. Californicus* (McGregor) female adults fed on *T. cinnabarinus* treated with sublethal concentrations of SYP-9625. (A) Control, (B) LC_10_, (C) LC_30_.

**Fig 10.**
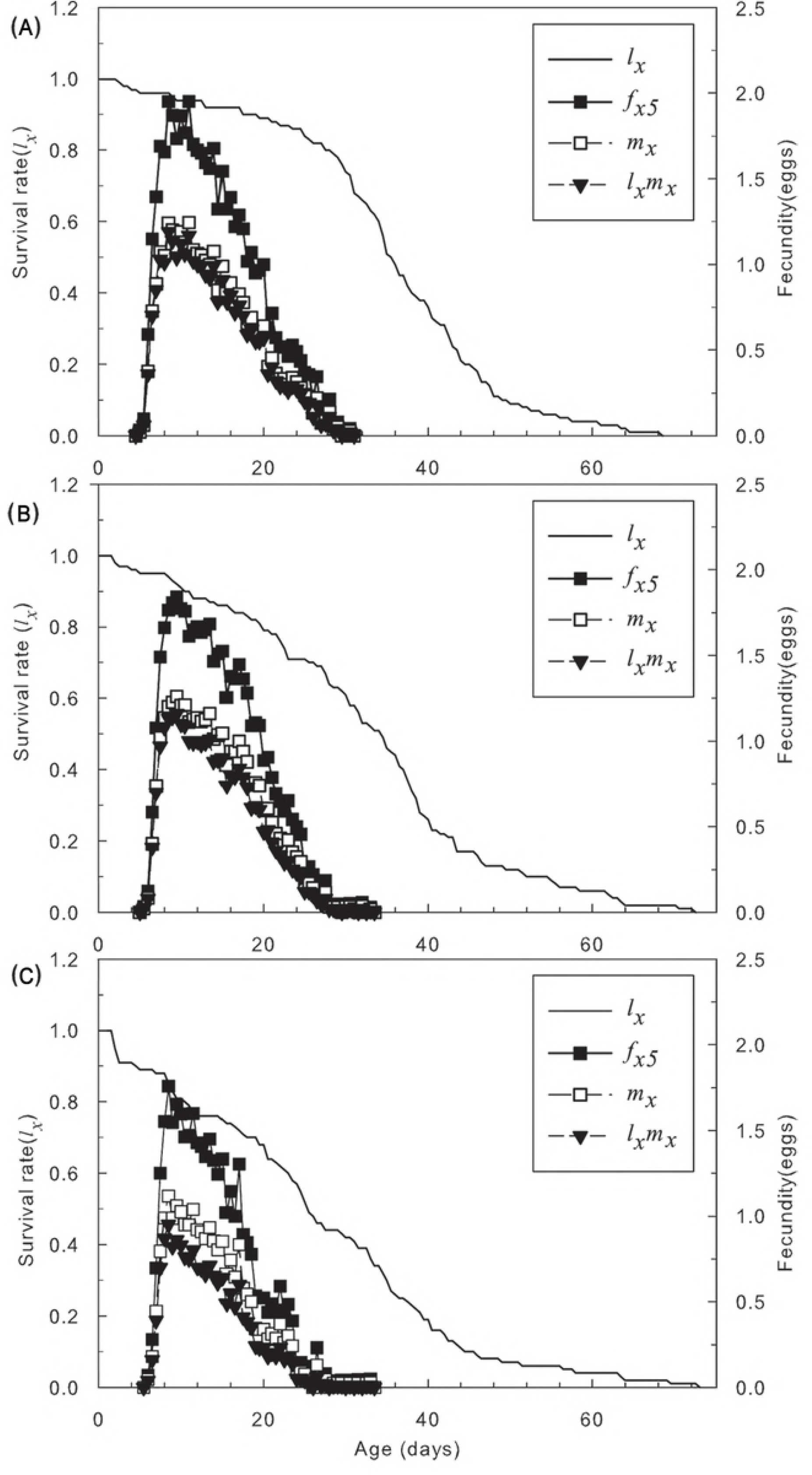
Age-specific survival rate (*l_x_*), female age-specific fecundity (*f*_*x*5_), age-specific fecundity of total population (*m_x_*), and age-specific maternity (*l_x_m_x_*) of offspring from *N. Californicus* (McGregor) female adult fed on *T. cinnabarinus* treated with sublethal concentrations of SYP-9625. (A) Control, (B) LC_10_, (C) LC_30_.

**Table 10.** Population life table parameters of offspring from *Neoseiulus californicus* fed on sublethal treated *Tetranychus cinnabarinus*

| Parameter | Control | SYP-9625 |  |
| --- | --- | --- | --- |
|  |  | LC<sub>10</sub> | LC<sub>30</sub> |
| Intrinsic rate of increase rate, $r_m$ (d<sup>-1</sup>) | 0.289±0.009<sup>a</sup> | 0.278±0.008<sup>a</sup> | 0.243±0.009<sup>b</sup> |
| Finite rate of increase, $\lambda$ (d<sup>-1</sup>) | 1.335±0.012<sup>a</sup> | 1.319±0.010<sup>a</sup> | 1.275±0.012<sup>b</sup> |
| Net reproductive rate, $R_0$<br>(offspring/individual) | 28.71±2.51<sup>a</sup> | 28.02±2.37<sup>a</sup> | 18.13±1.85<sup>b</sup> |
| Mean generation time, $T$ (d) | 11.61±0.15<sup>a</sup> | 12.01±0.15<sup>a</sup> | 11.91±0.16<sup>a</sup> |

### Effects of the application concentration on predatory capacity of *N. californicus*

The predatory capacity of *N. californicus* had an intrinsic acceleration owing to inverse density-dependent effects. After female adults were treated with 100 mg/L SYP-9625, there was no significant effect on predatory capacity against eggs of *T. cinnabarinus*. Although predatory capacity against larvae did not have dramatic change when there were 10 to 20 larvae of preys per leaf, predatory capacity against larvae declined significantly when prey density was 30 to 50 per leaf. The predatory capacity of female adults against nymphs changed irregularly (Table 11).

Table11 was uploaded as supported information in submission system.

The functional response data of *N. californicus* fit reasonably well to a type-II functional response of the Holling model. Functional response model and parameters of *N. californicus* treated with the application concentration changed in various stages (Table 12). The application concentration led to an increase in handling time and in attack rate against eggs and larvae. SYP-9625 exposure only affected the handling time negatively. Compared with the control, maximum attack rates (*T*/*Th*) decreased significantly except for nymphs. *a*/*Th* against eggs and nymphs increased 27.39% and 74.54%, respectively. *a*/*Th* against larvae and adults decreased 19.71% and 18.98%, respectively.

Table12 was uploaded as supported information in submission system.

### Effects on predatory capacity of *N. californicus* fed on sublethal treated *T. cinnabarinus*

The functional response model parameters of *N. californicus* fed on *T. cinnabarinus* treated with the sublethal concentrations changed in various stages (Table 13). There was no significant effect on predatory capacity against eggs of *T. cinnabarinus* when there were 20 or more eggs on a leaf, but predatory capacity increased when there were fewer than 20 eggs on a leaf. There was no apparent difference in predatory capacity against larvae among the two treatments and the control, but predatory capacity at LC_10_ was significantly less than the control when there were 40 larvae per leaf. There was no major difference in predatory capacity against nymphs among all treatments and the control while there were 10 to 20 nymphs per leaf. When nymph density increased to 25 to 30 per leaf, the predatory capacity of treatments was higher than the control. The predatory capacity of *N. californicus* at LC_10_ and LC_30_ was significantly higher than the control when 30 nymphs were set per leaf. There was little difference in predatory capacity among all treatments and the control while 5 to 15 adults were set per leaf. When nymph density increased to 20 to 25 per leaf, predatory capacities in the treatments were lower than the control.

Table13 was uploaded as supported information in submission system.

It was demonstrated that the functional response model was changed slightly based on the parameters in Table14. The sublethal concentrations led to an increase in attack rate against all the stages compared with the control. The attack rates at LC_10_ and LC_30_ against adults were increased 344.64% and 176.71%, respectively. Handling time against all stages did not differ at any concentration except the handling time against adults was longer than the control. The growth rate of *a/Th* against adults had a maximum value at LC_10_. When the concentration reached LC_30_, the value of *a/Th* was still higher than the control but a fall back trend appeared.

Table14 was uploaded as supported information in submission system.

## Discussion

### Sublethal effect of SYP-9625 on *T. cinnabarinus*

Acaricide and natural predators of mites were used as a successful combination of chemical and biological controls for pest management. Many authors considered the direct contact toxicity of acaricide against natural enemies but neglected sublethal effects, which intensify as time progresses.

**Our results show that the sublethal concentration of SYP-9625 can control the increasing population of *T. cinnabarinus* effectively**. And the overall impact on females is larger than males in *T. cinnabarinus* offspring. The population life table parameters except mean generation time of offspring treated with sublethal concentrations decreased significantly as the concentration increased.

### Effects of SYP-9625 on *N. californicus*

After determining the effect of the application concentration (100μg/mL) on *N. californicus* eggs, preadult duration, longevity and total life span of the treated *N. californicus* were mainly not varied. Other indexes including female proportion and adult emergence rate had less difference with the control as well. **All the results showed there was little influence on development and fecundity of *N. californicus* eggs. It was proved that the egg duration of *N. californicus* has good stress resistance to the application concentration of SYP-9625**.

**It indicates that exposure to the application concentration of SYP-9625 scarcely affected the survivorship of *N. californicus* and its subsequent generation**. *r_m_, λ* and *R_0_* in the offspring from females of *N. californicus* fed on *T. cinnabarinus* treated at LC_10_ were nearly same as the control, but these indices treated at LC_30_ were significantly reduced, which is consistent with part of previous findings [7]. The differences were potentially due to the various acaricide groups or concentrations and the different geographical populations of *N. californicus*.

### Effects of SYP-9625 on functional response of *N. californicus*

Natural enemies were not only exposed to sublethal concentration of insecticides but also fed on prey that exposed to sublethal concentration of insecticides. **No matter how predatory mites were treated with sublethal concentrations, no variation in the model of functional response was observed**. After *T. cinnabarinus* were treated with a sublethal concentration of SYP-9625, the attack rate of *N. californicus* against different instars of *T. cinnabarinus* increased, and a maximal increase slope showed in the adult treatment. Handling time of *N. californicus* against *T. cinnabarinus* treated with two sublethal concentrations (LC_10_, LC_30_) was longer than in the control. Furthermore, *a/Th* of treatments was higher than the control except for the larvae. Predation capacity of *N. californicus* against *T. cinnabarinus* treated with SYP-9625 was enhanced significantly, which is consistent with other results [4].

**Additionally, we found that there was no significant effect on predatory capacity of *N. californicus* against *T. cinnabarinus* eggs after it was treated with the SYP-9625**. The predatory capacity of *N. californicus* that treated with a higher dose against larvae and adults was significantly lower than the control. The attack rates against eggs and larvae were higher than the control, which is consistent with other research [29]. What counts is that we found the handling time of *N. californicus* was longer after treated with SYP-9625.

We suspect that the reason for this is that the toxicity of SYP-9625 against *N. californicus* is weak, and the Hormesis effect appeared at lower concentrations to stimulate the trophic behavior of *N. californicus*. The decline in functional response of larvae and adults was expected to display a normal range of change. It has been reported that the Hormesis effect at a low dose exists in various ecological populations, such as the predatory capacity of *Pardosa agrestis* treated with eight herbicides and *Supputius cincticeps* treated with sublethal concentrations of permethrin were evaluated, respectively [30, 31]. An alternative explanation could be that the application concentration of SYP-9625 stimulated *N. californicus* to take initiative to catch prey more often, but limited hunting behavior and other activity. Therefore, *N. californicus* was weaker to catch active life stages of prey such as adults than immobile life stages such as eggs. In conclusion, sublethal concentrations of SYP-9625 will partially weaken predation capacity of *N. californicus* against active *T. cinnabarinus* instars but will also enhance predation capacity against immobile instars.

It was found that a dose-dependent manner was inevitable, meaning that the sublethal effect of SYP-9625 was increased with increasing concentration. We maintain that lower concentration (LC_10_) of SYP-9625 is better for *N. californicus*. Scientific choice of miticides at reasonable concentrations can help optimal use of chemical and biological pest controls.

## Conclusions

We successfully assessed sublethal effects of SYP-9625 on *T. cinnabarinus*, effects of application concentration of SYP-9625 on the predatory mite *N. californicus* and functional responses of *N. californicus*. This study concludes that SYP-9625, especially lower concentration (LC_10_=0.375 μg/mL) can control the increasing population of *T. cinnabarinus* effectively and protect predatory mites appropriately. Additionally, we found that there was no significant effect on predatory capacity of *N. californicus* fed on treated *T. cinnabarinus* eggs. We suggest that the new acaricide SYP-9625 can be used with the release of predator mite *N. californicus*.

## References

1. Dejan M, Pantelija P, Slobodan M. Acaricides-biological profiles, effects and used in modern crop protection 2011. 39–62 p.

2. Yu H, Cheng Y, Xu M, Song Y, Luo Y, Li B. Synthesis, Acaricidal Activity and Structure-Activity Relationships of Pyrazolyl Acrylonitrile Derivatives. Journal of Agricultural & Food Chemistry. 2016; 51.

3. Alinejad M, Kheradmand K, Fathipour Y. Sublethal effects of fenazaquin on life table parameters of the predatory mite Amblyseius swirskii (Acari: Phytoseiidae). Exp Appl Acarol. 2014; 3: 361–373.

4. Poletti M, Maia A, Omoto C. Toxicity of neonicotinoid insecticides to Neoseiulus californicus and Phytoseiulus macropilis (Acari: Phytoseiidae) and their impact on functional response to Tetranychus urticae (Acari: Tetranychidae). Biological Control. 2007; 1: 30–36.

5. Lima DB, Monteiro VB, Guedes RNC, Siqueira HAA, Pallini A, Jr MGCG. Acaricide toxicity and synergism of fenpyroximate to the coconut mite predator Neoseiulus baraki. Biocontrol. 2013; 5: 595–605.

6. Park JJ, Kim M, Lee JH, Shin KI, Lee SE, Kim JG, et al. Sublethal effects of fenpyroximate and pyridaben on two predatory mite species, Neoseiulus womersleyi and Phytoseiulus persimilis (Acari, Phytoseiidae). Exp Appl Acarol. 2011; 3: 243–259.

7. Sá AP, Banyuls N, Santiago S, Mollá Ó, Jacas JA, Urbaneja A. Compatibility of Phytoseiulus persimilis and Neoseiulus californicus (Acari: Phytoseiidae) with imidacloprid to manage clementine nursery pests. Crop Protection. 2013; 1: 175–182.

8. Marafeli PP, Reis PR, Silveira ECd, Souza-Pimentel GC, Toledo MAd. Life history of Neoseiulus californicus (McGregor, 1954) (Acari: Phytoseiidae) fed with castor bean (Ricinus communisL.) pollen in laboratory conditions. Brazilian Journal of Biology. 2014; 3: 691–697. doi: 10.1590/bjb.2014.0079

9. Canlas LJ, Amano H, Ochiai N, TakedaM. Biology and predation of the Japanese strain of Neoseiulus californicus (McGregor)(Acari: Phytoseiidae). Systematic & Applied Acarology. 2012; 141: 141–157.

10. Fraulo AB, Liburd OE. Biological control of twospotted spider mite, Tetranychus urticae, with predatory mite, Neoseiulus californicus, in strawberries. Exp Appl Acarol. 2007; 2: 109–119.

11. Li DX, Tian J, Shen ZR. Functional response of the predator Scolothrips takahashii to hawthorn spider mite, Tetranychus viennensis: effect of age and temperature. Biocontrol. 2007; 1: 41–61.

12. Yorulmazsalman S, Ay R. Determination of the inheritance, cross-resistance and detoxifying enzyme levels of a laboratory-selected, spiromesifen-resistant population of the predatory mite Neoseiulus californicus (Acari: Phytoseiidae). Pest Manag Sci. 2013; 5: 819–826.

13. Li B, Yu H, Zhang H, Cheng Y, Luo Y, Wang L, et al. Pyrazolyl acrylonitrile compounds and uses thereof.: US; 2013.

14. Wang S, Tang X, Wang L, Zhang Y, Wu Q, Xie W. Effects of sublethal concentrations of bifenthrin on the two-spotted spider mite, Tetranychus urticae (Acari: Tetranychidae). Systematic & Applied Acarology. 1971; 4: 481–490.

15. Wang L, Zhang Y, Xie W, Wu Q, Wang S. Sublethal effects of spinetoram on the two-spotted spider mite, Tetranychus urticae (Acari: Tetranychidae). Pesticide Biochemistry & Physiology. 2016;: 102.

16. Carlo D, Valeria M, Alberto P, Marisa C, Marialivia L, Sauro S. Comparative toxicity of botanical and reduced-risk insecticides to Mediterranean populations of Tetranychus urticae and Phytoseiulus persimilis (Acari Tetranychidae, Phytoseiidae). Biological Control. 2008; 1: 16–21.

17. Li Q, Cui Q, Jiang C, Wang H, Yang Q, University SA. Control efficacy of Chinese Neoseiulus californicus(McGregor)population on Tetranychus cinnabarinus(Boisduval). Acta Phytophylacica Sinica. 2014; 3: 257–262.

18. Huang YB, Chi H. Life tables of Bactrocera cucurbitae (Diptera: Tephritidae): with an invalidation of the jackknife technique. Journal of Applied Entomology. 2013; 5: 327–339.

19. Kustutan O, Çakmak I. Development, fecundity, and prey consumption of Neoseiulus californicus (McGregor) fed Tetranychus cinnabarinus Boisduval. Turkish Journal of Agriculture & Forestry. 2014; 1: 19–28.

20. Williams FM, Juliano SA. FURTHER DIFFICULTIES IN THE ANALYSIS OF FUNCTIONAL-RESPONSE EXPERIMENTS AND A RESOLUTION. Canadian Entomologist. 1985; 5: 631–640.

21. Chi H. Life-Table Analysis Incorporating Both Sexes and Variable Development Rates Among Individuals. Environmental Entomology. 1988; 1: 26–34.

22. Chi H. Timing of control based on the stage structure of pest populations: a simulation approach. Journal of Economic Entomology. 1990; 4: 1143–1150.

23. Chi H, Liu H. Two new methods for study of insect population ecology. IEEE. 1985;:.

24. Cloyd RA, Galle CL, Keith SR. Compatibility of Three Miticides with the Predatory Mites Neoseiulus californicus McGregor and Phytoseiulus persimilis Athias-Henriot (Acari: Phytoseiidae). Hortscience A Publication of the American Society for Horticultural Science. 2006; 3: 707–710.

25. Lopez L, Smith HA, Hoy MA, Bloomquist JR. Acute Toxicity and Sublethal Effects of Fenpyroximate to Amblyseius swirskii (Acari: Phytoseiidae). Journal of Economic Entomology. 2015; 3: 1047–1053.

26. Ochiai N, Mizuno M, Mimori N, Miyake T, Dekeyser M, Canlas LJ, et al. Toxicity of bifenazate and its principal active metabolite, diazene, to Tetranychus urticae and Panonychus citri and their relative toxicity to the predaceous mites, Phytoseiulus persimilis and Neoseiulus californicus. Exp Appl Acarol. 2007; 3: 181–197.

27. Chi H. TWOSEX-MSChart: a computer program for the age-stage, two-sex life table analysis. 2012. http://140.120.197.173/Ecology/. National Chung Hsing University, Taichung Taiwan

28. Goodman D. Optimal Life Histories, Optimal Notation, and the Value of Reproductive Value. American Naturalist. 1982; 6: 803–823.

29. Martinou AF, Stavrinides MC. Effects of Sublethal Concentrations of Insecticides on the Functional Response of Two Mirid Generalist Predators. Plos One. 2015; 12: e0144413.

30. Korenko S, Niedobová J, Kolářová M, Hamouzová K, Kysilková K, Michalko R. The effect of eight common herbicides on the predatory activity of the agrobiont spider Pardosa agrestis. Biocontrol. 2016; 5: 1–11.

31. Zanuncio TV, Serrão JE, Zanuncio JC, Guedes RNC. Permethrin-induced hormesis on the predator Supputius cincticeps (Stål, 1860) (Heteroptera: Pentatomidae). Crop Protection. 2003; 7: 941–947.

